# Is the interaction of technology useful in laboratory haematology diagnostics?

**DOI:** 10.1101/2022.07.17.500333

**Authors:** Alessandra Falda, Marco Falda, Aurelio Pacioni, Giada Borgo, Rosolino Russelli, Antonio Antico

## Abstract

**Background:** Monoclonal B lymphocytosis (MBL) increases with age and individuals with high count MBL progress to chronic lymphocytic leukaemia requiring therapy at a rate of ∼1%-5% per year. These cases usually have atypical lymphocytes at the microscope, abnormal representation in the scattergram, and positivity of flags. Using XN9000 (Sysmex), we noticed cases of MBL without this correlation. We studied customized gates for discovering MBL cases of our interest.

**Methods:** We considered 212 peripheral blood samples with known phenotypes: 76.7% negative and 23.3% positive for B, T, or NK lymphocytes clones. We created gates studying the XN9000 FCS files in Diva software to identify new areas for better delimiting subpopulations of our interest and calculating sensitivity and specificity.

**Results:** We found significant differences between negative and positive groups for Q-flag “Blasts/Abn Lympho?” (B/AL) and LY-X (p <0.05) with lymphocyte counts below 5×10^9^/L.

A new gate P1 normalized by P2 (P1n) differentiated between phenotypes much better than Q-flag B/AL with lymphocyte counts ≤ 5 ×10^9^/L. Moreover, cases with MBL CD5 positive had higher medians (p <0.05).

**Conclusion:** We propose a gate P1n as a new Q-flag for lymphocytes count ≤ 5 ×10^9^/L, in order to hypothesize the presence of MBL CD5 positives.

## 1 INTRODUCTION

Nowadays, haematological instrumentation is very advanced and allows a great workflow, guaranteeing high quality in data analysis. Every instrument has a unit that processes and displays data generated by machines and software that allows viewing of parameters, graphics, and rules, and provides validation by the operator [1], after being expertly filtered.

Over the years, the technology has advanced enormously by refining algorithms that take into account the absolute values of the haematological series and the instrumental parameters [2] [3] [4] [5] [6] [7] [8] [9].

We know that microscopic review is still required whenever an instrumental anomaly (numerical, graphic, or usually triggered) occurs. Very experienced operators are needed, because the morphology is complex and also because, otherwise, there is a high degree of variability between observers.

Digital microscopy is increasingly used in diagnostic laboratories, even though the devices have a large footprint, are relatively expensive, and have limited slip parameters that must be adhered to generate accurate differentials [10] [11].

Flow cytometry gives the maximum guarantee of results by using specific markers in the recognition of abnormal cells or abnormal phenotypes. It is a second-level analysis and therefore it is not present in all laboratories.

Our laboratories are equipped with flow cytometry (FACSCanto II, Beckton-Dickinson) and XN9000 instrumentation (Sysmex, Japan). The parameters of XN9000 are expressed in arbitrary light scattering units (channels): LY-X describes the lateral scatter signal and depends on the cellular complexity (SSC), LY-Y represents the lateral fluorescence light intensity (SFL), and LY-Z is related to cell volume (FSC). The white blood cell channel (WDF) distinguishes all leukocyte populations except basophils.

In our experience, we observed some cases where there was no correspondence between the presence of abnormal lymphocytes on microscopic examination, an abnormal representation of the lymphocytes in the WDF scattergram, and the presence of the Q flag “Blasts/Abn Lympho?” (B/AL). These cases were then confirmed in the cytometric study as clonal B lymphocytosis.

When clonal B lymphocytes are less than 5×10^9^/L, in the absence of lymphadenopathy and/or organomegaly, these cases are defined as monoclonal B lymphocytosis (MBL) [12]. MBL becomes more common with age, and individuals with high-count MBL develop chronic lymphocytic leukaemia (CLL) that requires therapy at a rate of 1%-5% per year [13] [14] [15].

In 2021, Cornet et al. observed significant differences between neoplastic and reactive lymphocytosis for LY-X and LY-Z (Instruments XN-1000, Sysmex), but not for Q-flag B/AL and “Atypical Lympho?” [16]. They focused their attention on lymphocyte counts above 5×10^9^/L, because over this threshold it is still mandatory to investigate the case in-depth to differentiate between secondary and neoplastic origin [17] [18]. They observed that in the latter scenario, most cases had an LY-X value greater than 78.5 and an LY-Z value greater than 58.0, independently of the instrumental XN flags [17].

We explored the use of instrument parameters and XN9000 flags to characterize the differences between neoplastic and non-neoplastic lymphocytosis, dividing the cases between “positive” and “negative” and then keeping only the lymphocytosis less than 5×10^9^/L. Another aim was to focus on the profile XN-DIFF (WDF channel) for differentiating lymphocytes: we tried to identify new gates in flow cytometry standard (FCS) files of XN studied with the Diva software (Becton-Dickinson FACS Canto II flow cytometer) to identify cell populations of interest.

## 2 MATERIALS AND METHODS

Three hundred and eighty-three patients were considered from the routine workflow of the Clinical Pathology Department of ULSS7 Pedemontana (Vicenza, Italy).

Among patients with two samples taken at different times (one for blood count and one for lymphocyte typing), we retained samples with a reference change value for lymphocyte count lower than 28% [19].

After this step, the dataset was composed of 288 cases: 184 with a single blood sample and 104 with two different withdrawals.

Peripheral blood samples were collected in K3-EDTA anticoagulated vacutainer (Beckton Dickinson, San Jose, California, USA). The peripheral phenotype and analysis on the Sysmex XN-9000 series haematology analyzer were known for all cases.

Since the XN instrumentation with the WPC channel was only present in one of our multicenter laboratories, our attention was placed on the Q-flag B/AL and not on the single flag after reflecting on the WPC channel.

Two commercial control samples (XN-Check, level 1 and 2; Sysmex) were measured daily and sent to the Sysmex web service (Caresphere™ XQC, Sysmex); manufacturer-based standard operating procedures were followed. This was done with six machines from two locations. Results of both internal controls met predefined performance indicators. The analyzer uses fluorescence flow cytometry for the differential count of leukocytes and adds further information about cell sizes (forward diffusion; FSC), internal structure/granularity (lateral dispersion; SSC), and DNA/RNA content (intensity fluorescence; SFL).

We considered the phenotype from the 4^th^ edition of WHO Classification for categorizing B lymphocyte clones on peripheral blood [12].

Among 123 positive patients, 58 were clonal B lymphocytes “CLL”, 13 “atypical CLL” and 40 “not CLL”. Twelve patients had T or NK neoplasm.

One hundred and sixty-five were negative cases, represented by 71 healthy (normal distribution of T, B and NK lymphocytes) and 94 reactive cases (cases with quantitative alteration of T, B or NK, but without phenotypic anomalies).

The percentage of males and females was 52.4% and 47.6%, respectively; the age was higher in the patient group (median 75 years vs 57). The ratio between negative and positive cases was 1.293. There was a higher frequency of “positive” samples in males (63.5%).

In the context of lymphocyte counts ≤ 5×10^9^/L (212 patients), we imported the FCS files of the WDF scattergram from XN9000 to Canto II (BD) to better study cases with atypical lymphocytes in the morphology of peripheral blood, with known phenotype and no Q-flag B/AL flagged. We designed 3 gates: P3 to represent the total area of interest, P1 and P2 for studying the areas of our particular interest (Figure 1).

**Figure 1.**
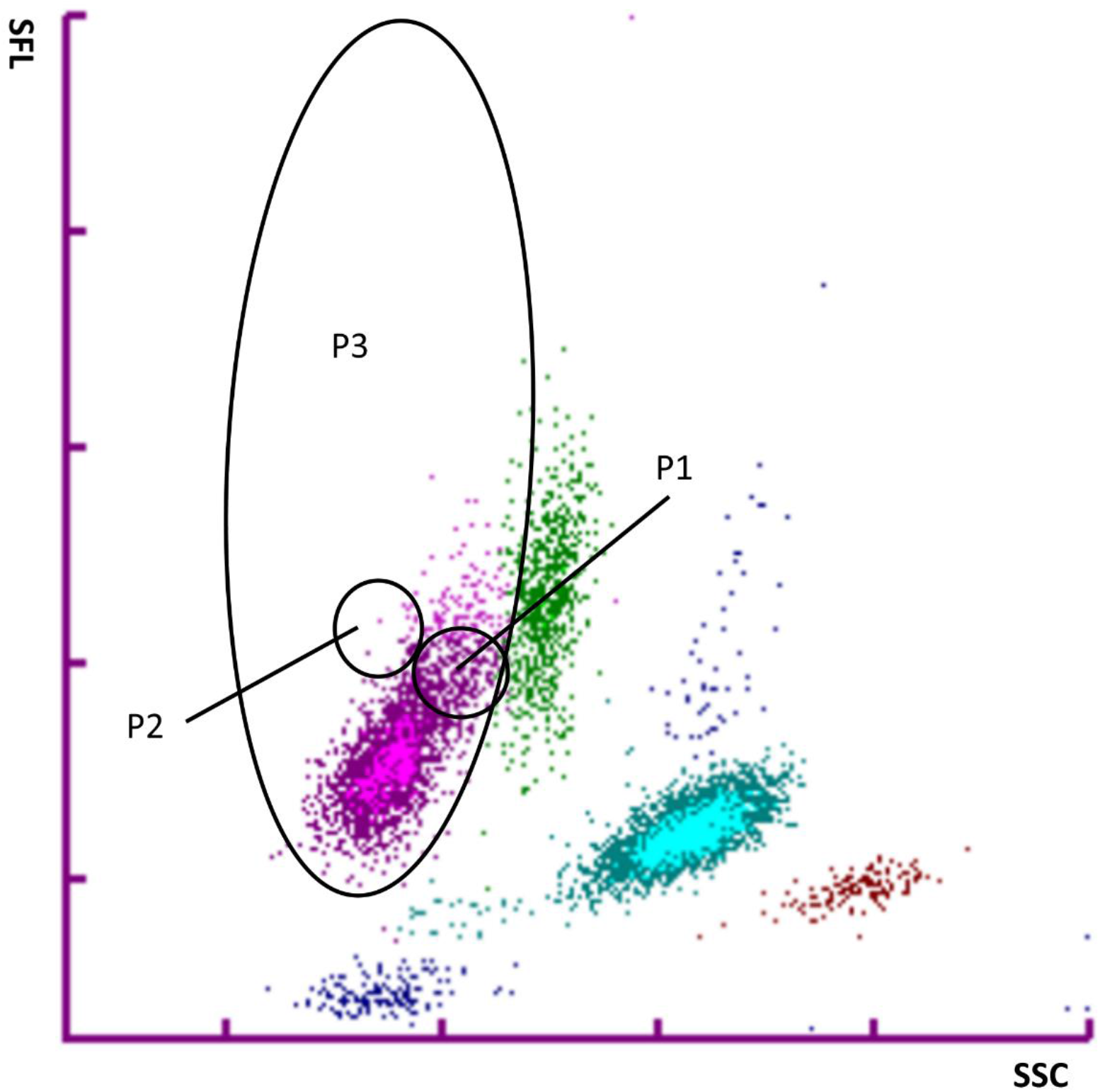
Personalized gates: P3 represents the total area of interest, P2 and P1 are for studying the areas of our interest.

P1 gate represents an area with channels for SSC and SFL slightly above the normal lymphocytes for XN. To better focus on lymphocyte phenotypes, we decided to exclude P3 and consider only P1/P2, where P2 allowed us to “normalize” P1 (gate P1n in the following).

In this last analysis, we did not consider T and NK haematological neoplasms, because there were only 6 cases with lymphocyte counts ≤ 5×10^9^/L.

With lymphocyte count ≤ 5×10^9^/L we observed: CLL median 0.422 B lymphocytes ×10^9^/L, atypical CLL 0.965 ×10^9^/L, “not CLL” 0.305 B lymphocytes ×10^9^/L, healthy 0.164 ×10^9^/L, healthy + polyclonal B 0.191 ×10^9^/L, reactive 0.079 ×10^9^/L and reactive + polyclonal B 0.131 ×10^9^/L.

## Statistical analysis

The statistical analysis was performed using GNU R 4.1.0 [20]. A p-value less than 0.05 was considered statistically significant.

We used a two-tailed Mann-Whitney test between negative and positive cases for the evaluation of custom gates on lymphocyte counts ≤ 5×10^9^/L.

Kruskal-Wallis test with a 95% confidence interval was used to study differences among cytometric groups. Conover-Iman test was used for *post hoc* comparisons.

## 3 RESULTS

For lymphocytes counts ≤ 5×10^9^/L we found a significant p-value for LY-X (p = 0.0086) between negative and positive groups: positive cases had lymphocytes with a significantly higher SSC. For SFL we didn’t find significant differences (p = 0.0525), as well as in the evaluation of LY-Z (p = 0.4790). When counts were greater than 5×10^9^/L, groups were imbalanced and so we did not apply statistical tests.

In the context of lymphocyte counts ≤ 5×10^9^/L, to better study the areas of our interest in the lymphocytes gate, we imported the FCS files of the WDF scattergram from XN9000 to Canto II: in a first analysis, we observed statistically significant differences for the Q-flag B/AL between negative and positive cases, up to lymphocytes counts of 4.0×10^9^/L and with an AUC of 0.632. The Q-flag had good specificity, while sensitivity decreased under 50%. Using the P1% parent (P1/P3), statistically significant differences were seen between negative and positive cases for counts up to 2.0×10^9^/L. Sensitivity was better than the Q-flag (Table 1).

**Table 1.** Sensitivity and specificity of Q-flag B/AL and P1% parent to differentiate negative from positive groups (ALC= absolute lymphocytes count; AUC = area under curve).

| N. cases | ALC ($\times 10^9/L$) | p-value | Sensitivity (%) | Specificity (%) | AUC |
| --- | --- | --- | --- | --- | --- |
| **Q-flag B/AL** |  |  |  |  |  |
| 206 | $\leq 5.0$ | 0.007 | 0.542 | 0.785 | 0.627 |
| 180 | $\leq 4.0$ | 0.012 | 0.459 | 0.867 | 0.632 |
| 143 | $\leq 3.0$ | 0.485 | 0.440 | 0.797 | 0.544 |
| 100 | $\leq 2.0$ | 0.576 | 0.278 | 0.890 | 0.459 |
| 56 | $\leq 1.5$ | 0.351 | 0.308 | 0.837 | 0.418 |
| **P1% parent** |  |  |  |  |  |
| 206 | $\leq 5.0$ | 4.76E-08 | 0.729 | 0.772 | 0.760 |
| 180 | $\leq 4.0$ | 1.39E-05 | 0.703 | 0.762 | 0.732 |
| 143 | $\leq 3.0$ | 0.001 | 0.720 | 0.729 | 0.719 |
| 100 | $\leq 2.0$ | 0.014 | 0.722 | 0.707 | 0.686 |
| 56 | $\leq 1.5$ | 0.281 | 0.769 | 0.558 | 0.600 |

We expanded the analysis by evaluating these differences between the various cytometric groups. From Kruskal-Wallis analysis, both Q-flag B/AL and gate P1n showed a statistically significant difference (p<0.05) between at least one of the cytometric groups; this result was found for Q-flag B/AL up to lymphocyte counts of ≤ 4.0×10^9^/L, while for gate P1n up to counts ≤ 1.5×10^9^/L (Table 2).

**Table 2.**
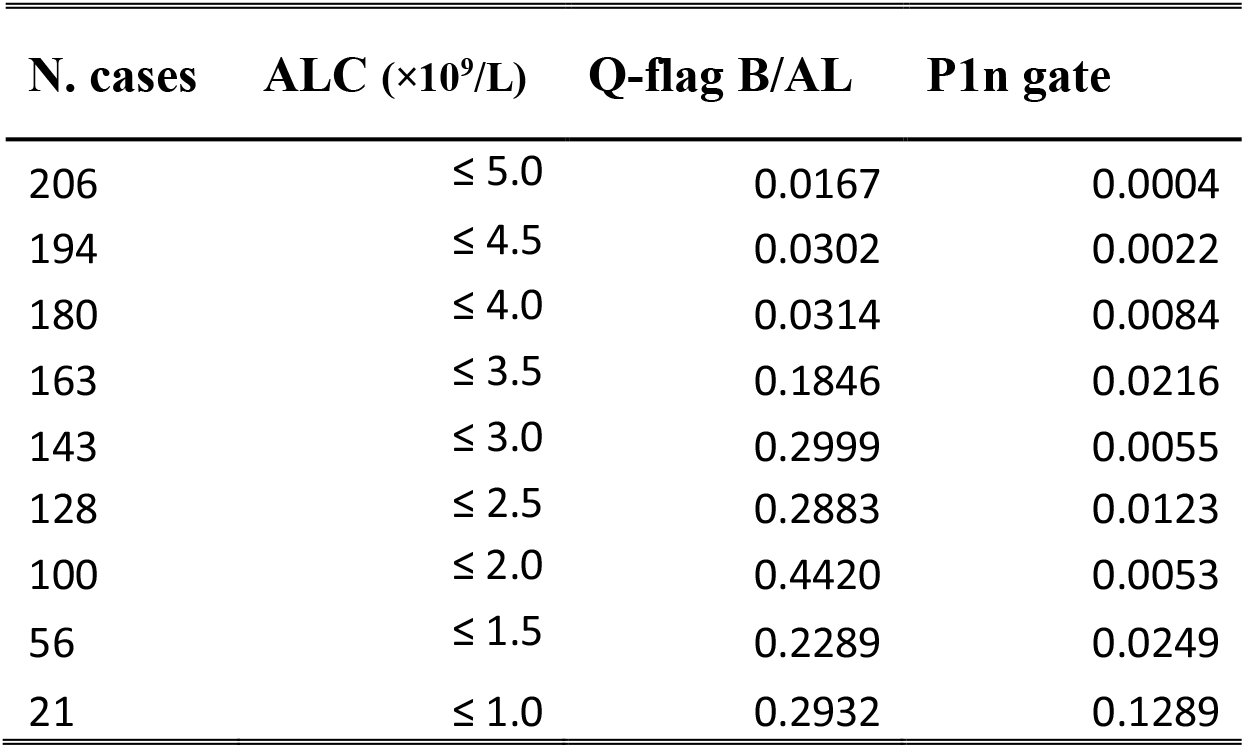
P-values of Q-flag B/AL and P1n gate in phenotypic groups when ALC is under 5 × 10^9^/L (ALC= absolute lymphocytes count).

When we considered medians of Q-flag B/AL, all cytometric groups had a value less than 100 (discriminant threshold for identifying positive cases) (Figure 2a).

**Figure 2.**
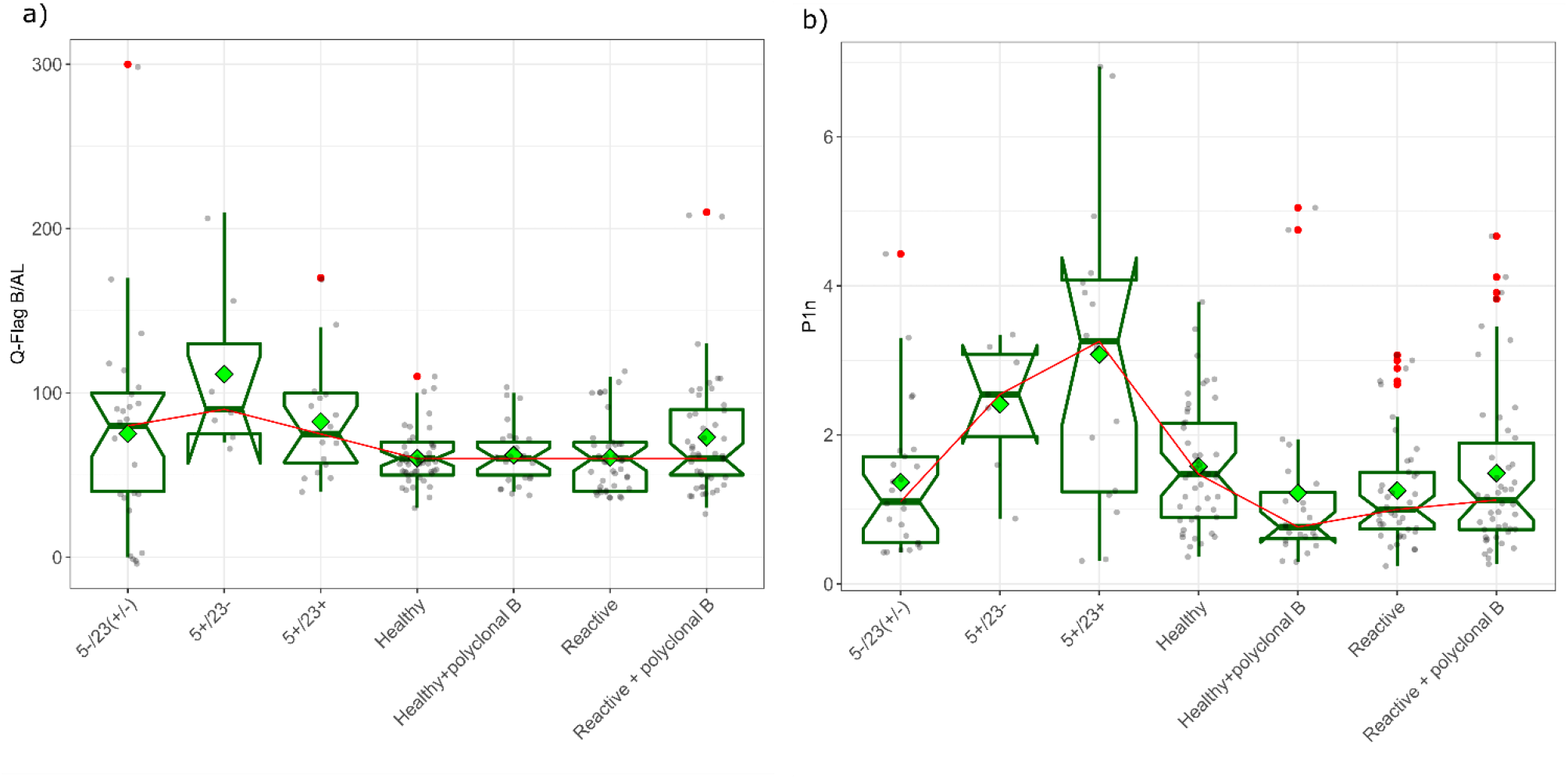
Boxplots of Q-flag B/AL (3a) and gate P1n (3b) by cytometric groups with lymphocyte counts ≤ 5×109/L. (Reactive: T and/or B and/or NK lymphocytes outside the range of normality, poly B: polyclonal B lymphocytes; green diamonds: means).

In the case of gate P1n, we observed that only CD5 positive clonal B cells had medians greater than 2 (Figure 2b), moreover by applying a Conover-Iman test we observed that those two cytometric groups (CD5+/CD23+ and CD5+/CD23±) had statistically significant differences with respect to the other groups (p <0.05) also with smaller lymphocytes counts (Table 3).

**Table 3.** Pairwise P1n differences between cytometric groups. Lymphocytes count ≤ 5 ×109/L. Reject H0 if p ≤ alpha/2 (**: p < 0.05; *: p < 0.10).

| **P.adjusted** | $\leq 5 \times 10^9/L$ | $\leq 4 \times 10^9/L$ | $\leq 3 \times 10^9/L$ | $\leq 2 \times 10^9/L$ | $\leq 1.5 \times 10^9/L$ |
| --- | --- | --- | --- | --- | --- |
| 5+/23- - Reactive | 0.047** | 0.188 | 0.247 | 0.177 | 0.333 |
| 5+/23+ - Reactive | 0.007** | 0.090* | 0.378 | 0.674 | 0.941 |
| 5+/23- - Reactive + polyc. B | 0.081* | 0.335 | 0.439 | 0.565 | 0.833 |
| 5+/23+ - Reactive + polyc. B | 0.021** | 0.241 | 0.674 | 0.655 | 1.000 |
| 5+/23- - Healthy + polyc. B | 0.010** | 0.045** | 0.041** | 0.063* | 0.038** |
| 5+/23+ - Healthy + polyc. B | 0.001** | 0.013** | 0.051* | 0.202 | 0.926 |

## 4 DISCUSSION

When performing a blood count, it is critical to recognize lymphocytic diseases and, when appropriate, to recommend a phenotypic study. The microscopic examination of cases can also be difficult for the operator because there may be morphological overlaps or co-occurrence of reactive and neoplastic lymphocyte patterns. As a result, in our study, we examined the FCS files of the WDF scattergram in 206 cases with known phenotypes to determine the clinical usefulness of various instrumental characteristics (absolute counts, flags, instrumental parameters) in screening cases with or without chronic lymphoproliferative disease in peripheral blood. We focused on lymphocyte counts ≤ 5×10^9^/L, where it is more common to diagnose MBL. We know that when the most sensitive analytical techniques have been used, the prevalence of MBL can reach 20% in subjects over the age of 60 and 75% in subjects over the age of 90 [16].

We studied the WDF scattergram of XN-9000 that represents the distribution of leukocytes based on two parameters: LY-X (SSC) and side fluorescence light (SFL). We found that LY-X was significantly higher in the neoplastic cases than in the others (p= 0.0086). For SFL, we didn’t find significant differences between positives and negatives (p=0.0525), as well as in the evaluation of LY-Z, a parameter that describes cell size (p=0.4790). To our knowledge, we are the first to have done this analysis on lymphocyte counts in the range of normality.

Regarding differences in graphical representations, Sale et al. in their 2016 work with XN-1000 demonstrated that in cases of chronic lymphocytic leukaemia (CLL), leukemic lymphocytes tend to cluster in the typical spherical lymphocyte cluster in the WDF scattergram [19]. We think that this is also possible for lymphocyte counts ≤ 5 ×10^9^/L, in the analysis with XN-9000 instrumentation.

In our experience, we observed that there is not always a correspondence between the presence of abnormal lymphocytes on morphological examination and the manifestation of instrumental anomalies. This also occurs in the presence of abnormal distribution of events in the lymphocyte population with a WDF scattergram, where the B/AL flag does not appear. For this reason, we analyzed XN FCS files with Diva software.

We stratified the positive and negative groups based on lymphocyte counts ≤ 5 × 10^9^/L. The sensitivity of the Q flag decreases with the lymphocyte count (below 50% to ≤ 4.0 × 10^9^/L) because several neoplastic lymphocytes fall below the detection threshold of the Q flag, hence the number of false-negative cases increases.

The sensitivity of the P1 gate tends to enhance when the lymphocyte counts decrease because its representation in neoplastic cases moves near the monocytes, and so the number of false negatives reduces. Conversely, false positives increase and specificity decreases.

We didn’t observe this behaviour in reactive cases, where an increase in SFL is prevalent.

When we considered the medians of Q-flag B/AL, all cytometric groups had a value less than 100 (discriminating threshold for identifying positive cases). We then evaluated the P1n gate in lymphocyte counts ≤ 5 × 10^9^/L: median of P1n events was higher in CD5 positive clonal B cases than in the other phenotypes, in one group even for lymphocyte counts up to ≤ 1.5 × 10^9^/L (p <0.05).

There were many more B lymphocytes per microliter in the clonal cases (CLL median 0.430 B lymphocytes ×10^9^/L, atypical CLL 0.872 ×10^9^/L, not CLL 0.305 B lymphocytes ×10^9^/L) compared to the negative group (healthy 0.164 ×10^9^/L, healthy + poly B 0.193 ×10^9^/L, delta 0.079 ×10^9^/L and delta + poly B 0.131 ×10^9^/L).

The absolute number of clonal B lymphocytes reported by flow cytometry is much greater than the number of events that fall inside the P1n gate drawn on the representation of the WDF scattergram of XN. For this reason, we hypothesize that the P1n gate is not the representation of the whole clone, but that inside gate P1n there are many more events related to B neoplastic CD5 positive cases with respect to negative cases. We are aware that this model needs to be validated with a larger cohort.

It is difficult to give a univocal interpretation of this result; however, in the context of neoplastic B lymphocytosis heterogeneity is observed both at a morphological and also phenotypic level. It is plausible that this is also reflected in the graphic representation of the single case. For low lymphocyte counts, in XN-9000 we observed that the atypical lymphocytes area protrudes towards the monocytes, which is our gate P1 (Figure 1). This may be due to the co-presence of more complex and less mature cells, such as in atypical/mixed CLL and some leukemic lymphomas.

Regarding differences in graphical representations, Sale et al. in their 2016 work with XN-1000 demonstrated that in cases of chronic lymphocytic leukaemia (CLL), leukemic lymphocytes tend to cluster in the typical spherical lymphocyte cluster in the WDF scattergram [20].

There are Authors that plotted gates on haematological software to study haematological malignancies, reactive lymphocytosis and malaria [22], but few studies utilise gating strategies applied in different software as in our case [19] [20].

We think it is possible to adopt gate P1n as a new Q-flag in XN instrumentation to suspect the presence of CD5 positive B clones in lymphocyte counts ≤ 5 × 10^9^/L. It could allow to reduce the operator-dependent evaluation and ease the recognition of neoplastic clones, even of small dimensions. We think that this might be possible because the units of measurement referred to the physical or fluorescence parameters are the same. We acknowledge that we have to verify this hypothesis because there are differences depending on manufacturers, models and maps, which are strictly instrumental dependent.

## AUTHOR CONTRIBUTIONS

A. Falda conceived the cytometric analysis and designed the gates, contributed to the data analysis, and wrote the manuscript. M. Falda realized the data analysis and helped with the manuscript preparation. A. Pacioni was involved in data analysis. G. Borgo and R. Russelli created the data library. A. Antico was involved in the manuscript preparation.

